# Design and validation of a new flat-sheet membrane bioreactor system for bioprocessing research requiring very low gas fluxes

**DOI:** 10.1101/2025.04.15.649005

**Authors:** Mei Zhou, Ibrahim Olanrewaju Bello, Jose Antonio Magdalena, Joseph G. Usack

**Author notes:** **Corresponding author: Joseph G. Usack, Ph.D.**, Department of Food Science & Technology, University of Georgia, 100 Cedar Street, Athens, GA 30602.

## Abstract

Precise and consistent gas dosing at trace levels remains a significant challenge in bioprocessing and biotechnology research. This study introduces a novel flat-sheet membrane bioreactor for controlled gas delivery without bubble formation. The system relies on gas diffusion mediated by interchangeable membrane cassettes and incorporates mechanical stirring near the membrane surface to promote gas dispersal and mitigate biofouling spanning long operating periods. Comprehensive characterization of mass transfer properties using oxygen gas as a test case revealed that the volumetric mass transfer coefficient (K_L_a) remained stable under varying operational conditions, including gas partial pressure, back pressure, and total gas flow rate. The maximum oxygen flux under specific operational conditions flexibly ranged from 15.9E-03 ± 6.3E-03 mol·min^−1^ to 1.08 ± 0.17 mol·min^−1^ for various porous membranes and from 23.3E-03 ± 3.5E-03 mol·min^−1^ to 0.161 ± 0.044 mol·min^−1^ for the non-porous membranes, indicating the bioreactor can serve as an experimental platform for a variety of applications. A biological validation study using starch-containing wastewater demonstrated the feasibility of continuous microaerobic oxygen dosing, achieving reproducible performance and maintaining stable operation over 320 days. These findings highlight the potential of the developed system as a reliable experimental platform for investigating bioprocesses dependent on trace gas supply. Future improvements should focus on scaling up the system, addressing porous membrane wetting, and expanding its applicability to other gases and bioprocessing applications.

## 1. Introduction

Precise control of trace gas dosing in bioreactor systems is critical for studying key metabolic pathways, bioenergetics, and process tolerances [1, 2]. For example, hydrogen gas partial pressure (pH_2_) influences the thermodynamic favorability of fatty acid oxidation metabolism in anaerobic digestion (AD) at very low concentration ranges (*e.g.*, 1E-03 to 1E-05 atm) [1]. Moreover, many industrial gas fermentation processes, such as biomethanation, syngas fermentation, and CO_2_-to-X, are adversely affected by trace impurities in the feedstock gas (*e.g.*, oxygen (O_2_) and hydrogen sulfide (H_2_S)) [2, 3]. These impurities undermine process performance by causing 1) microbial toxicity, 2) enzyme inhibition, 3) alteration of electron flow, or 4) byproduct formation [2]. Microaeration, which involves adding very small amounts of oxygen to establish specific oxidation-reduction potentials (ORP) during fermentation, has emerged as a promising strategy for enhancing these bioprocesses. Applications include improving organic matter hydrolysis [4], mitigating recalcitrant pollutant degradation [5, 6], increasing methane yields in AD systems [7, 8], promoting H_2_S [9] and carboxylic acid oxidation [10, 11], and steering the bioprocess towards or away from specific end products, such as caproic acid [12] and caprylic acid [13].

However, conventional gas dosing methods make it challenging to quantify the amount of gas transferred into the liquid phase of biological systems. For instance, exogenous oxygen or air sparging has been the most common approach for microaeration, where oxygen or air is sparged into the bioreactor headspace [14], influent [5], or the liquid phase [15]. The gas sparging method presents several limitations: 1) it creates spatial gas concentration gradients within the bioreactor, leading to uneven oxygen distributions that stress obligate anaerobes and compromise bioprocess stability [16], and 2) not all gas bubbles dissolve during sparging, making it challenging to estimate or model biological oxygen availability. As a result, studies often report microaeration intensity values, given as the gas flow rate per unit reactor volume (*e.g.*, mL O_2_·L^−1^·d^−1^), to indirectly estimate biological oxygen availability, complicating cross-study comparisons [17]. Thus, there is a need to develop a more robust experimental platform with precise and quantifiable gas dosing capabilities that researchers can use to characterize the effects of gas composition and gas availability on bioprocess performance in various applications.

Gas transfer membranes, with well-defined gas diffusion rates, offer a potential solution to more precisely and uniformly dose gases at trace levels in bioreactor systems. Gas transfer membranes are typically used to supply gases for bioprocesses at high dosing rates. One example is aerating the bioreactor using submerged flow-through hollow fiber polydimethyl siloxane (PDMS) membranes to achieve oxygen transfer rates as high as 35 g O_2_·m^−2^·day^−1^ [18]. Dead-end hollow fiber membranes are another typical gas-supply configuration in membrane bioreactors [19, 20].

However, these membrane configurations are susceptible to non-uniform gas transfer, pore blocking, and membrane fouling over time, which reduce gas transfer efficiency and increase the risk of membrane rupture due to high-pressure build-up [21]. Rotating membranes were developed to increase the shear stress on the membrane surface, but fouling is still problematic [22]. Progressive fouling precludes research requiring precise and consistent gas dosing over long study periods. Still, membrane bioreactors have the potential to be exact and consistent, provided the applied gas pressure remains below the bubble-point pressure of the membrane and technical issues such as pore blocking and fouling are addressed [7]. Under these conditions, diffusion through the membrane matrix controls gas transfer into the bioreactor, which can be readily quantified via direct abiotic measurement or modeling. Furthermore, because the membrane matrix is physically isolated from the bioreactor broth under non-wetting conditions, gas transfer rates would be minimally affected by biological changes occurring in the bioreactor, remaining consistent during operation.

This study introduces and evaluates a novel diffusion-driven flat-sheet membrane bioreactor for experimental applications requiring precise gas dosing at trace levels for extended operating periods. The system integrates mechanical stirring near the membrane surface to improve gas dispersion and reduce biofouling, promoting uniform gas dosing. The primary objectives of this research are to 1) characterize the gas transfer performance of the bioreactor under varying membrane types and operational conditions and 2) validate its utility in providing consistent gas dosing during extended operation. Oxygen gas was used to characterize bioreactor gas transfer performance, and a long-term biological microaeration study was used to validate its gas dosing consistency. The study aims to advance gas transfer technology for experimental and practical bioprocessing applications.

## 2. Materials and methods

### 2.1 Membrane bioreactor design and reactor configurations

Several membrane bioreactor prototypes were built and tested for functionality. The final flat-sheet membrane bioreactor design (**Fig. 1**) comprised the following components: 1) a gas supply system that delivered dynamic gas mixtures using mass flow controllers (MFCs) (FS-201CV-1K0-RBD-00-K, Bronkhorst, Kamen, Germany); 2) a bioreactor vessel with an open bottom that included a temperature-controlled water bath, a magnetic stirring plate, a jacketed glass reactor vessel, an pH/ORP probe, and a removable head plate; 3) a gas flow-through chamber with a surficial orifice, and an interchangeable membrane cassette; 4) an optical trace oxygen detection module; and 5) off-gas lines connected to back-pressure regulators and airlocks.

**Fig. 1.**
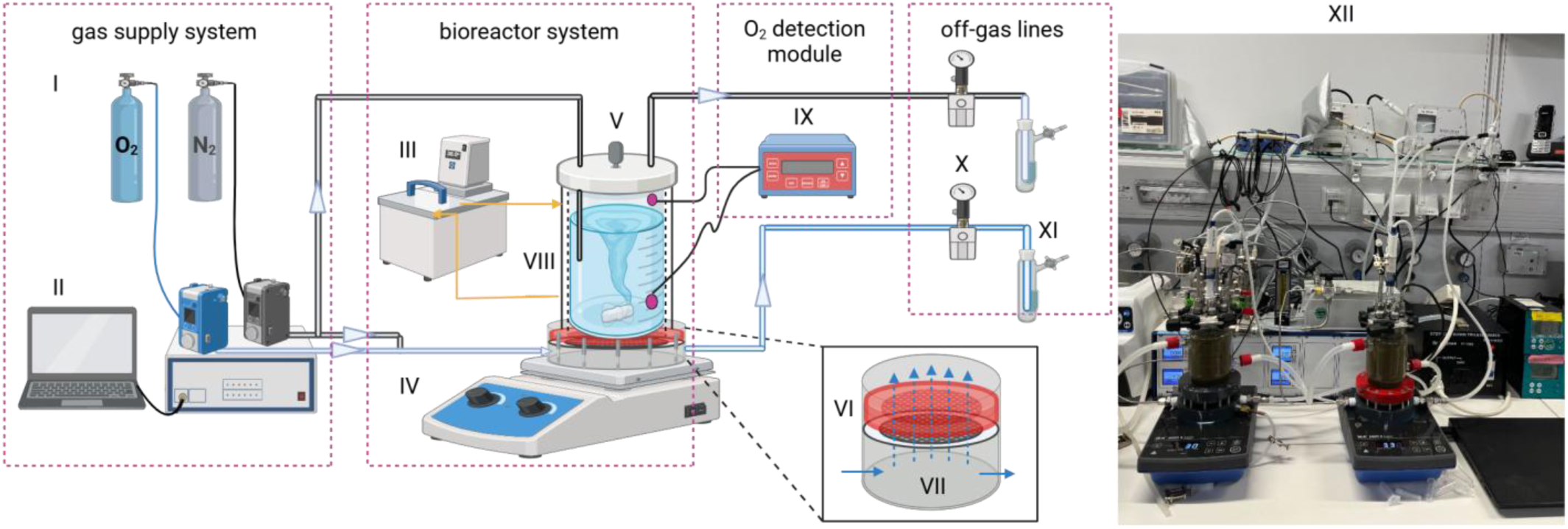
Schematic diagram of the bioreactor system. I: Oxygen (O_2_) and nitrogen (N_2_) tanks, II: Gas supply system comprising mass flow controllers, a gas mixer, and gas flow software, III: Temperature-controlled circulating water bath, IV: Magnetic stirring plate, V: Jacketed glass bioreactor vessel, VI: Interchangeable membrane cassette, VII: Gas flow-through chamber, VIII: Optical non-invasive O_2_ sensor spots, IX: Gas or dissolved O_2_ concentration sensor, X: Back-pressure regulators, XI: Airlock, XII: Picture of the membrane bioreactor system. Created with BioRender.com.

FlowDDE V 4.83 and FlowView 7 V 1.23 software (Bronkhorst, Kamen, Germany) controlled the mixing of oxygen and nitrogen and regulated mixed gas flow to the gas flow-through chamber. Here, gas permeates the flat membrane, dissolving into the reactor broth proportionate to the applied gas partial pressure. Note: all gas flow rates (*i.e.*, L·h^−1^) in this work have been normalized to standard conditions (*i.e.*, T = 20℃, P = 101 kPa). The pressure meter monitored the pressure of the mixed gas flow (EL-PRESS, Bronkhorst, Kamen, Germany), and a differential pressure meter (CR410, Ehdis, Dongguan, China) located between the off-gas lines of the gas side and the bioreactor headspace measured the differential pressure across the membrane. The interchangeable sealed membrane cassette provided mechanical support for the flat-sheet membrane at elevated gas pressures. The interchangeable membrane cassette comprised two 3-mm rubber gaskets (NR/SBR, 40 +/− 5 Shore A, Reiff Technische Produkte GmbH, Reutlingen, Germany), the interior of which held fritted glass discs (60mm, filter discs, biplane, series 16, porosity 0, VitraPOR®, ROBU Glasfilter, Hattert, Germany). The membrane was cut to size and then compression-held between the top and bottom gasket-glass disc assembly, creating a hermetic seal with the membrane.

The stirring plate (KMO 3 basic, IKA, Staufen, Germany) and an internal magnetic stir bar at the membrane cassette surface were designed to mitigate biofouling and promote gas dispersion. The combined thickness of the flow-through gas chamber and membrane cassette was constrained to 15 cm, not to exceed the working range of the magnetic stir plate. Finally, a trace oxygen detection system (OXY-4 SMA, PreSens, Regensburg, Germany) and non-invasive optical oxygen sensors spots (SP-PSt6-NAU-D5-YOP of 0 – 2 mg·L^−1^ and SP-PSt3-NAU of 0 – 45 mg·L^−1^, Model, PreSens, Regensburg, Germany) were used to continuously measure and log oxygen concentrations in the liquid and gaseous phases with the software of PreSens Measurement Studio 2 Version 4.0.0 (PreSens, Regensburg, Germany).

A computational fluid dynamics (CFD) simulation was performed to characterize fluid mixing behavior within the bioreactor based on mixing speeds using OpenFOAM (**Section SI l**). This open-source software uses a finite-volume method to solve partial differential equations. The mixing phenomenon was simulated using the incompressible VoF solver – a solver that handles transient, incompressible two-phase flows. The CFD analysis indicated the bioreactor was well-mixed with relatively uniform fluid velocities within the liquid volume for each simulated mixing speed (*i.e.,* 250 rpm, 350 rpm, and 450 rpm) with no significant dead zones (**Fig. S1**).

### 2.2. Abiotic tests evaluating gas transfer properties of the designed membrane bioreactor

#### 2.2.1 Experimental design and materials

Abiotic tests focused on characterizing the bioreactor’s precision, dynamic oxygen flux range, and parameter sensitivity using the volumetric mass transfer coefficient (K_L_a) and oxygen flux as metrics. The operating parameters such as membrane type (*e.g.*, material, porosity, thickness), oxygen partial pressure (%), gas flow back pressure (bar), rotational speed of the stir bar (rpm), and total gas flow rate (L·h^−1^) were systematically varied to define the bioreactor’s capabilities at a constant temperature setting (*i.e.,* 37°C) (**Table 1**). Various commercially available sheet membranes were utilized, including porous polyvinylidene fluoride (PVDF), polypropylene (PP), and polytetrafluorethylene (PTFE), and non-porous silicone membranes with different thicknesses (**Table 2**). Gas pressures at the membrane were set by regulating the back-pressure regulator at the outlet of the gas flow-through chamber. New membranes were used for each trial.

**Table 1.** Experimental design used to evaluate the effect of operational parameters on the gas transfer properties of the membrane bioreactor.

| Tests | Membrane | Total Gas<br>Flow Rate<br>(L·h<sup>-1</sup>) | Oxygen Partial<br>Pressure (%) | Reactor<br>Temperature<br>(°C) | Rotational<br>Speed<br>(rpm) | Gas Back<br>pressure<br>(bar) |
| --- | --- | --- | --- | --- | --- | --- |
| No.1 | PVDF<sup>a</sup> | 63 | 5.6, 21, 44.4,<br>66.7, 100 | 37 | 250 | 0 |
|  | PP<sup>a</sup> | 63 | 3.0, 21, 44.4,<br>66.7, 100 | 37 | 250 | 0 |
|  | PTFE<sup>a</sup> | 63 | 10, 21, 33.3,<br>44.4, 100 | 37 | 250 | 0 |
|  | Silicone 1 | 63 | 7.8, 21, 33.3,<br>66.7, 100 | 37 | 250 | 0 |
| No.2 | PVDF<sup>a</sup> | 7 | 100 | 25 | 250 | 0.05, 0.1,<br>0.50, 0.75, 1 |
| No.3 | PVDF<sup>a</sup> | 7 | 100 | 25 | 150, 250, 350,<br>450 | 0.4 |

<sup>a</sup> Polyvinylidene fluoride (PVDF); Polypropylene (PP); Polytetrafluoroethylene (PTFE)

**Table 2.** Characteristics of the membranes used to evaluate the effect of membrane type on the gas transfer properties of the bioreactor.

| Membrane | PTFE<sup>a</sup> | PP<sup>a</sup> | PVDF<sup>a</sup> | Silicone 1 | Silicone 2 |
| --- | --- | --- | --- | --- | --- |
| Manufacturer<br>(serial number) | BOLA<br>(N1617-04) | Dorsan<br>(M090PP020) | Carl-Roth<br>(2831.1) | KT<br>(0505773) | Modular<br>(0227347) |
| Pore size (µm) | 0.2 | 0.2 | 0.2 | - | - |
| Thickness (µm) | 200 | 1447.8 | 125.0 ± 25 | 300 | 3000 |
| Specific surface<br>area (dm<sup>2</sup>) | 28.27 | 28.27 | 28.27 | 28.27 | 28.27 |
| Bubble point (bar) | - | 0.8 | - | - | - |

#### 2.1.2 Calculations

The oxygen mass balance from the gas flow-through chamber to the bioreactor vessel was independently validated using abiotic tests with anaerobic distilled water for each membrane and operational condition. The oxygen mass balance in the well-mixed liquid phase was determined with the following equations [23]:

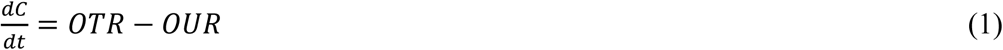

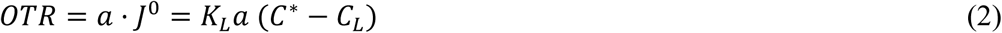

Where *dC/dt* is the oxygen concentration differential per unit time; *OTR* is the oxygen transfer rate from the gas to the liquid phase; *OUR* is the oxygen uptake rate by the microorganisms, which is zero for the abiotic tests; *J^0^*is the molar flux of oxygen through the gas-liquid interface; *a* is the gas-liquid interfacial area per unit of liquid volume; *K_L_* is the overall mass transfer coefficient; *C^⁎^* is the oxygen saturation concentration in the bulk liquid in equilibrium with the bulk gas phase; *C_L_* is the dissolved oxygen (DO) concentration in the bulk liquid at a specific time.

Since measuring *K_L_* and *a* separately is complex, the transport of oxygen from gas to liquid is characterized by the volumetric mass transfer coefficient (*K_L_a*) based on the dynamic change of DO concentration in the bioreactor vessel over time. The dynamic change in the DO concentration between two different times and oxygen flux is expressed as:

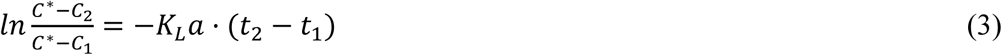

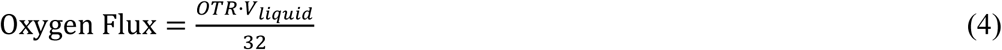

Where *C_2_* is the DO concentration at *t_2_* time, *C_1_* and *t_1_* are the concentration and time at the beginning of each test.

### 2.2. Biotic microaeration study testing system stability during long-term operation

#### 2.2.1 Experimental conditions

The biotic microaeration case study utilized flat-sheet membrane Silicone 2 (3 mm thick) in two identical bioreactors operated at 37°C with continuous stirring at 250 rpm. The linear fit shows that the maximum oxygen flux through the membrane is proportional to the oxygen partial pressure. Each bioreactor had a fill volume of 200 mL, with batch-fed influent and effluent volumes of 10 mL per day, corresponding to a hydraulic retention time (HRT) of 20 days and an organic loading rate (OLR) of 2 g COD·L^−1^·day^−1^. The inoculum and the composition of the feeding mixture are detailed in the Supplementary Information (**Section SI Ⅱ**). During the startup phase, both bioreactors received a supply of 16.8 L·h^−1^ N_2_ to the gas flow-through chamber with the back-pressure regulators fully open. On Day 157, the mass flow controllers for the experimental microaeration bioreactor (R1-MA) and the control anaerobic digestion bioreactor (R2-AD) were calibrated to 16.31 L·h⁻¹ and 16.75 L·h⁻¹, respectively, resulting in roughly equal gas flow rates of 210.15 mL·min⁻¹ for R1-MA and 210.91 mL·min⁻¹ for R2-AD at standard conditions (*i.e.*, 0 ℃ and 1 atm) using the same gas meter (µFlow, BPC Instruments, Lund, Sweden). In subsequent experimental periods, the oxygen partial pressure in R1-MA was adjusted by substituting nitrogen gas with air, based on bioreactor performance, while maintaining consistent total gas flow rates.

#### 2.2.2 Analytical methods

Volumetric biogas production rates were measured by the µFlow gas meters and automatically normalized to 0℃ and 1 atm conditions with the built-in sensors. Daily volumetric biogas production rates were multiplied with the mid-day methane content in the biogas to estimate daily methane production. Biogas composition was sampled at 13.5 h after feeding with a 500 μL gas-tight syringe (Hamilton, Reno, USA) and analyzed using two gas chromatographs (GCs) (SRI 8610C; SRI Instruments, Torrance, CA), modified from the method described by Usack *et al.* [24]. The GC for O_2_ and N_2_ measurement was equipped with a molsieve 13x column (length 3 m, outer diameter 1/8”, SRI Instruments, Torrance, CA) and a thermal conductivity detector (TCD) with argon gas as the carrier gas. The GC oven was operated at isothermal conditions (*i.e.*, 52℃). The other GC for H_2_, CO_2_, and CH_4_ measurement was equipped with a HaySep-D packed Teflon column (length 3 m, outer diameter 1/8”, SRI Instruments, Torrance, CA) and a TCD and a flame ionization detector (FID) with N_2_ as the carrier gas. The GC oven was operated at isothermal conditions (*i.e.,* 70℃).

Carboxylic acid concentrations were determined using an Agilent 7890B GC (Agilent Technologies Inc., Santa Clara, CA, USA) equipped with a capillary column (DB Fatwax ultra inert, 30 m × 0.25 mm × 0.25 µm; Agilent Technologies Inc., Santa Clara, CA) and an FID detector with hydrogen gas as mobile phase, according to Gemeinhardt *et al.* [13]. Analysis of total/soluble chemical oxygen demand (tCOD/sCOD), total solids (TS), volatile solids (VS), and volatile suspended solids (VSS) was performed according to Standard Methods [25]. Starch concentration was analyzed using the total starch HK assay kit (K-TSHK, Megazyme, Th. Geyer GmbH & Co. KG, Renningen, Germany). Ammonium concentrations were measured on Day 195 following standard methods DIN 38406/ISO 11732. The filtered (0.2μm) samples reacted with salicylate and dichloroisocyanuric acid, forming a blue complex that was measured at 660 nm. pH of liquid samples was measured periodically *ex-situ* using the ceramic pH probe (12×120 mm, pHenomenal® 221, VWR International GmbH, Darmstadt, Germany). A redox micro ORP electrode (Mettler-Toledo 51343203, Mettler-Toledo InLab, Gießen, Germany) was connected through a dedicated port in the bioreactor lid of R1-MA to measure ORP *in-situ*.

## 3 Results and discussions

### 3.1 Gas transfer properties of the membrane bioreactor

#### 3.1.1 Effects of gas composition, gas flow rate, and membrane type on oxygen transfer properties (No.1)

This section evaluates the oxygen transfer properties of different membranes by monitoring the dynamic change in DO concentration over time in the bioreactor vessel. Saturation concentrations at each condition were simulated using the correlation between DO concentration and time (**Fig. 2a**). The dynamic change in DO concentration over time generated a linear relationship when plotted using **Eq. 3**, and K_L_a was extracted from the slope of the line (**Fig. 2b**). The OTR and oxygen flux decreased over time due to the declining gradients of C* and C_L_ (**Fig. 2c**). In addition to the membrane type, the oxygen transfer properties of specific membrane cassette configurations were determined by modifying oxygen partial pressure in the gas flow.

**Fig. 2.**
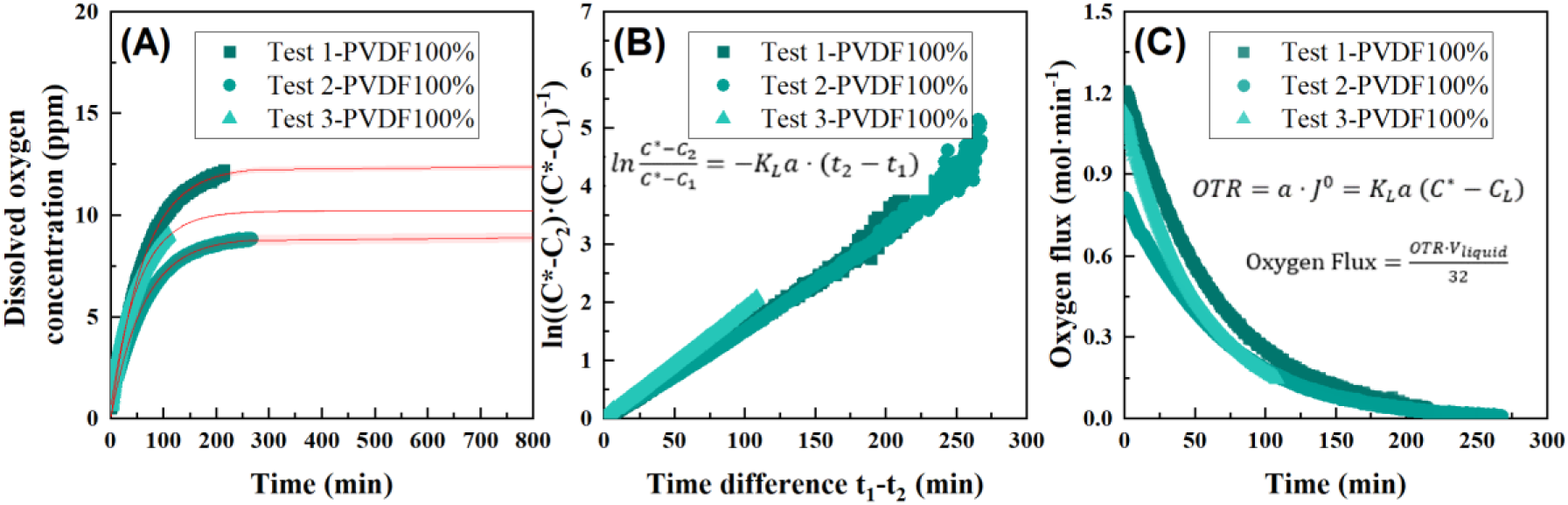
Example triplicate plots for determining the oxygen mass transfer coefficient (K_L_a) of the PVDF membranes with oxygen partial pressure set to 100%: (A) Modeled saturation concentration (C*) (red lines) from the time course of the measured (scatters) dissolved oxygen concentration in the bioreactor vessel; (B) K_L_a extraction from the slope of the line of the calculated ln((C*-C_2_)·(C*-C_1_)^−1^) and (t_1_-t_2_) (b); (C) Time course of the oxygen flux through the membrane.

We first determined the gas transfer properties of the designed bench-scale membrane bioreactor, with a liquid volume of 200 mL, using various hydrophobic membranes at 37℃ and 250 rpm. During these trials, no back pressure was applied to the gas flow-through chamber or bioreactor, resulting in approximately equal total pressure on both sides of the membrane. Here, gas transport was driven mainly by diffusion based on the oxygen partial pressure differential across the membrane. The oxygen partial pressure differential was set by the N_2_:O_2_ gas composition passing through the gas flow-through chamber, assuming an oxygen partial pressure of zero in the bioreactor, which contained anaerobic water.

The results indicated that increasing oxygen partial pressure did not significantly affect K_L_a because the slope of linear-fitted trend lines was not significantly different from zero (p > 0.05, **Fig. 3A**). This suggests that gas-liquid transfer at the membrane surface did not strongly constrain the overall flux. In contrast, the membrane type remarkedly affected K_L_a. The estimated K_L_a of the non-porous silicone membrane (*i.e.*, 7.13E-03 ± 3.29E-04 min^−1^) was considerably lower than that of the porous polymer membranes (*i.e.*, 14.1E-03 ± 4.01E-04 min^−1^ of PTFE, 16.3E-03 ± 4.22E-04 min^−1^ of PP, 18.1E-03 ± 4.22E-06 min^−1^ of PVDF) (p < 0.0001 in **Fig. 3A**). Notably, PVDF exhibited the highest K_L_a, followed by PP and PTFE. Porous membranes (*i.e.*, PVDF, PP, and PTFE) showed higher OTRs than the non-porous silicone membrane due to their higher permeability and lower resistance to diffusion. These findings are consistent with the results of Luqmani *et al.* [26], who demonstrated higher CO_2_ flux by the porous hollow fiber membrane contactor than the one containing non-porous material.

**Fig. 3.**
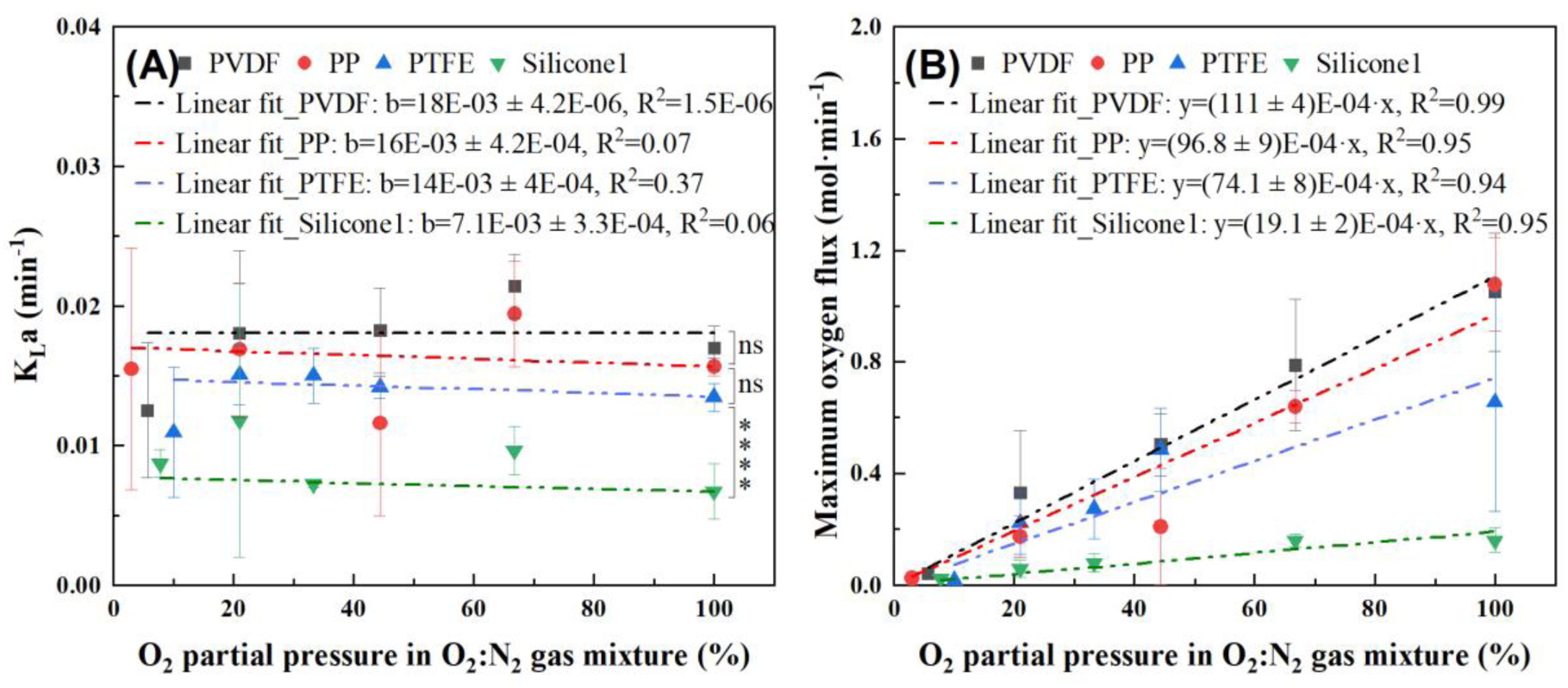
(A) The measured (scatters) and simulated (lines, slopes are 0) volumetric mass transfer coefficient (*K_L_a*); (B) The effect of oxygen partial pressure on the measured maximum oxygen flux (scatters) and the linear fit (lines). Various membrane types were tested: porous membranes (porosity = 0.2 μm) including Polyvinylidene fluoride (PVDF), Polypropylene (PP), and Polytetrafluoroethylene (PTFE), and a non-porous membrane (*i.e.*, Silicone 1). We assessed the significance between *K_L_a* of different membranes using Two-Sample t-tests (p-value with asterisks in the figure, ns is no significant difference, and **** indicates < 0.0001).

While the oxygen partial pressure differential does not significantly affect the K_L_a for each membrane, it does result in directly proportional changes in the oxygen flux (**Fig. 3B**). When the oxygen partial pressure was increased from 3% to 100%, the maximum oxygen flux through porous membranes ranged from 15.9 E-03 ± 6.3 E-03 mol·min^−1^ to 1.08 ± 0.17 mol·min^−1^. The non-porous silicone membrane provided a narrower flux range from 23.3E-03 ± 3.5E-03 mol·min^−1^ to 0.161 ± 0.044 mol·min^−1^ with oxygen pressures between 7.8% and 100%. These results demonstrate that the membrane bioreactor can be custom-configured with different membrane types and gas partial pressures to achieve different dosing ranges for trace gas dosing applications. The porous membrane types offer wider dosing ranges but are less precise than the non-porous membrane. The oxygen flux range can be increased by applying positive back pressure in the gas flow-through chamber. While oxygen was used as the model gas in this case study, similar functionality and responses can be expected for other gases possessing similar solubilities (*e.g.*, H_2_, CH_4_, CO).

In addition to the oxygen partial pressure and membrane types, we also evaluated the effect of membrane thickness and total gas flow rates on gas transfer properties. A thicker 3-mm membrane (*i.e.,* Silicone 2) was tested in subsequent trials. Unlike the thin membranes, the 3-mm-thick silicone membrane did not require additional structural support using fritted glass discs within the membrane cassette. Replicate measurements indicated that removing the fritted glass discs reduced the variation in K_L_a measurements, with the relative standard deviation (RSD) decreasing from 43% to 4% (ratio of standard deviation and mean of K_L_a of the first two rows in **Table 3**). This finding suggests that the variability in preliminary K_L_a measurements was primarily due to the presence of the fritted glass support discs. We selected fritted glass discs with pore sizes much larger than the membrane pores, assuming this would minimize gas hold-up. However, the fritted glass discs had this very effect, perhaps due to their high pore volume, tortuosity, and irregular (pore) shape. Future designs should apply a support material with a similarly large but more regular pore structure.

**Table 3.** Effect of removing the glass disc and the total gas flow rate on the gas transfer properties of the bioreactor using a 3-mm-thick silicone membrane.

| Glass discs | Total Gas Flow<br>Rate ($L \cdot h^{-1}$) | Oxygen Partial<br>Pressure (%) | $K_{La}$ | Maximum Oxygen Flux<br>($mol \cdot min^{-1}$) |
| --- | --- | --- | --- | --- |
| with | 63 | 80 | $0.0082 \pm 0.0035$ | $0.0666 \pm 0.0408$ |
| without | 63 | 80 | $0.0134 \pm 0.0006$ | $0.4562 \pm 0.0268$ |
| without | 16.8 | 100 | 0.0110 | 0.0391 |
| without | 63 | 100 | 0.0115 | 0.4522 |

Hence, because of its higher precision, we used the 3-mm-thick silicone membrane to investigate the effect of total gas flow rate on oxygen transfer properties. The results showed that while the total flow rate of the O₂:N₂ mixture (100%:0%) (last two rows in **Table 3**) did not significantly impact the K_L_a, it did alter the oxygen flux through the membrane by an order of magnitude. These results indicate that the total gas flow rate can be used as an additional control parameter to adjust the oxygen transfer rate in the membrane bioreactor.

#### 3.1.2 Gas transfer rates can be further adjusted by applying back-pressure or varying mixing speeds (No.2 – 3)

In addition to membrane type and oxygen partial pressure on the gas side, the gas transfer rates can be further adjusted by altering specific operational parameters, such as the applied back pressures and rotational speed of the magnetic stir bar. Preliminary tests showed that the system with PVDF could withstand up to 1 bar of back pressure. Back pressures higher than 1 bar caused membrane deformation despite being structurally supported with glass discs on both sides. Additionally, back pressures higher than 0.6 bar led to gas bubble formation. Therefore, for all tests in this section, a near-equal back pressure in the bioreactor headspace was applied to avoid membrane deformation and bubble formation.

In this set of tests, pure oxygen at 7 L·h^−1^ was supplied to the gas flow-through chamber containing a PVDF membrane cassette. Back pressure was systematically varied. Results indicated that the back pressure did not significantly affect the K_L_a, as the slope of the trend line was not significantly different from zero (p > 0.05, black line in **Fig. 4A**). However, it did have a direct effect on the oxygen saturation concentration, which increased proportionally with the back pressure (blue scatters in **Fig. 4A**). The maximum oxygen flux responded similarly, exhibiting a linear response ranging from 0.23 mol·min^−1^ to 4.31 mol·min^−1^ for a back pressure range of 0 to 1 bar, respectively (pink scatters in **Fig. 4A**). Higher back pressures increase the gas flux by effectively increasing the O_2_ partial pressure and, subsequently, the gas saturation concentration (C*). Adjusting the back pressure from 0 to 1 bar increased the oxygen flux by approximately one order of magnitude, considerably extending the functional dosing range of the membrane bioreactor system.

**Fig. 4.**
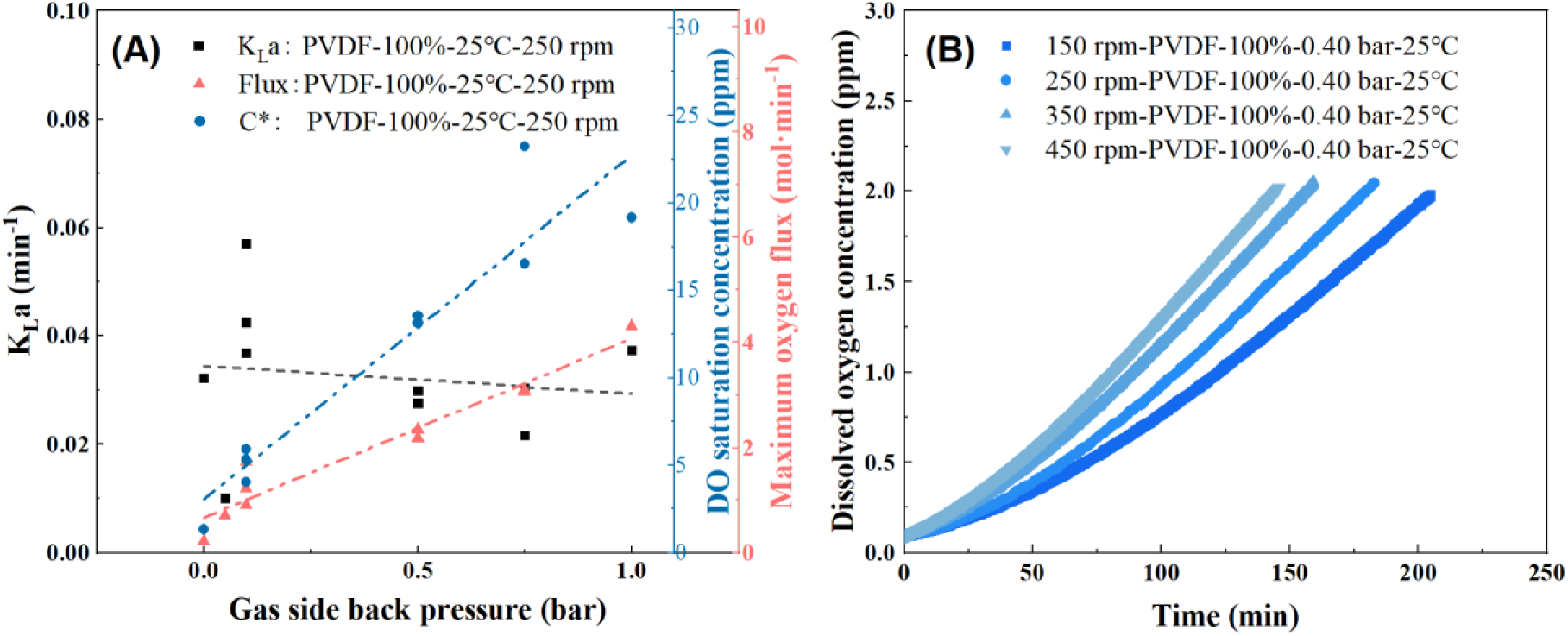
(A) The measured (scatters) and modeled (lines) volumetric mass transfer coefficient (*K_L_a*), dissolved oxygen (DO) saturation concentration (C*), and the maximum oxygen flux of the Polyvinylidene fluoride (PVDF) membrane cassette at different back pressures; (B) The effect of the rotational speed (rpm) of the magnetic stir bar on the DO concentration versus time.

The rotational speed of the magnetic stir bar on the membrane surface can also be adjusted to change the gas transfer rates. The rotational speed changes the turbulence level and contact area between the gas and liquid phases. The DO time course trends varied with rotational speed and were highest at 450 rpm, followed by 350 rpm, 250 rpm, and 150 rpm (**Fig. 4B**). The saturation concentration of oxygen was not modeled for the tested speeds, but faster speeds are expected to reduce the equilibration time during oxygen dosing while promoting spatial uniformity in the bioreactor. The rotational speed also affects the shear stress at the membrane surface, which could help mitigate membrane fouling. Shear stress generally decreases membrane fouling [27], but the relationship is complex and depends on the application. For example, higher shear stress helps dislodge or prevent the attachment of biofouling agents; however, certain microbes (*e.g.*, *Pseudomonas aeruginosa*) react to shear stress by producing more extracellular polymeric substances, leading to more strongly adhering biofilms [28].

#### 3.1.3 Technical challenges and uncertainty

The membrane bioreactor was designed to use hydrophobic membranes. The porous membranes are still susceptible to partial or complete wetting despite being hydrophobic. After 24 hours of operation, water penetrated each porous membrane and accumulated inside the gas flow-through chamber. The PTFE membrane exhibited the highest membrane wettability (*i.e.,* shortest time to wetting), followed by the PP and PVDF membranes. The pore size of each membrane was 0.2 μm, and the operating conditions were the same in all trials (*i.e.,* 25°C, 250 rpm). Water penetrates the membrane pores because of their low surface tensions. Membrane wetting decreases mass transfer due to lower gas diffusivity in the wetted membrane phase [29]. Thus, new strategies are needed to solve the problem of porous membrane wetting, such as developing superhydrophobic membrane materials. What is worth noting is that the non-porous membrane showed superior resistance to wetting. Additionally, investigating the long-term stability and performance under continuous operation will be essential to ensure the membrane bioreactor can be reliably implemented in practical applications. Future research should focus on testing the membrane bioreactor design at larger scales and under more varied conditions to confirm its robustness.

### 3.2 Biological case study: microaeration treatment of starch-containing wastewater

#### 3.2.1 Microaeration flexibility, precision, and stability in a practical case

Two membrane bioreactors were operated for over 320 days using the 3-mm silicone membrane with no fritted glass disc. After resolving microbiome-related issues during the startup phase (detailed in the Supplementary Information) (**Section SI Ⅱ**), the bioreactors performed consistently at a 30-day HRT and an OLR of 1.3 g COD·L^−1^·day^−1^ from Day 160 onward. Between days 160 and 174, the observed stable DO concentration (0.009 ppm) and ORP values (– 508 ± 3 mV) were indicative of strict anaerobic conditions (data shown only for R1-MA in **Fig. 5A**). Additionally, the biogas composition was also similar (**Fig. 5B**). There were no significant differences (p < 0.05) in biogas and methane production rates between the two bioreactors (**Fig. 5C, 5D**). These findings demonstrate that the membrane bioreactor is gas-tight and suitable for anaerobic biological processes.

**Fig. 5.**
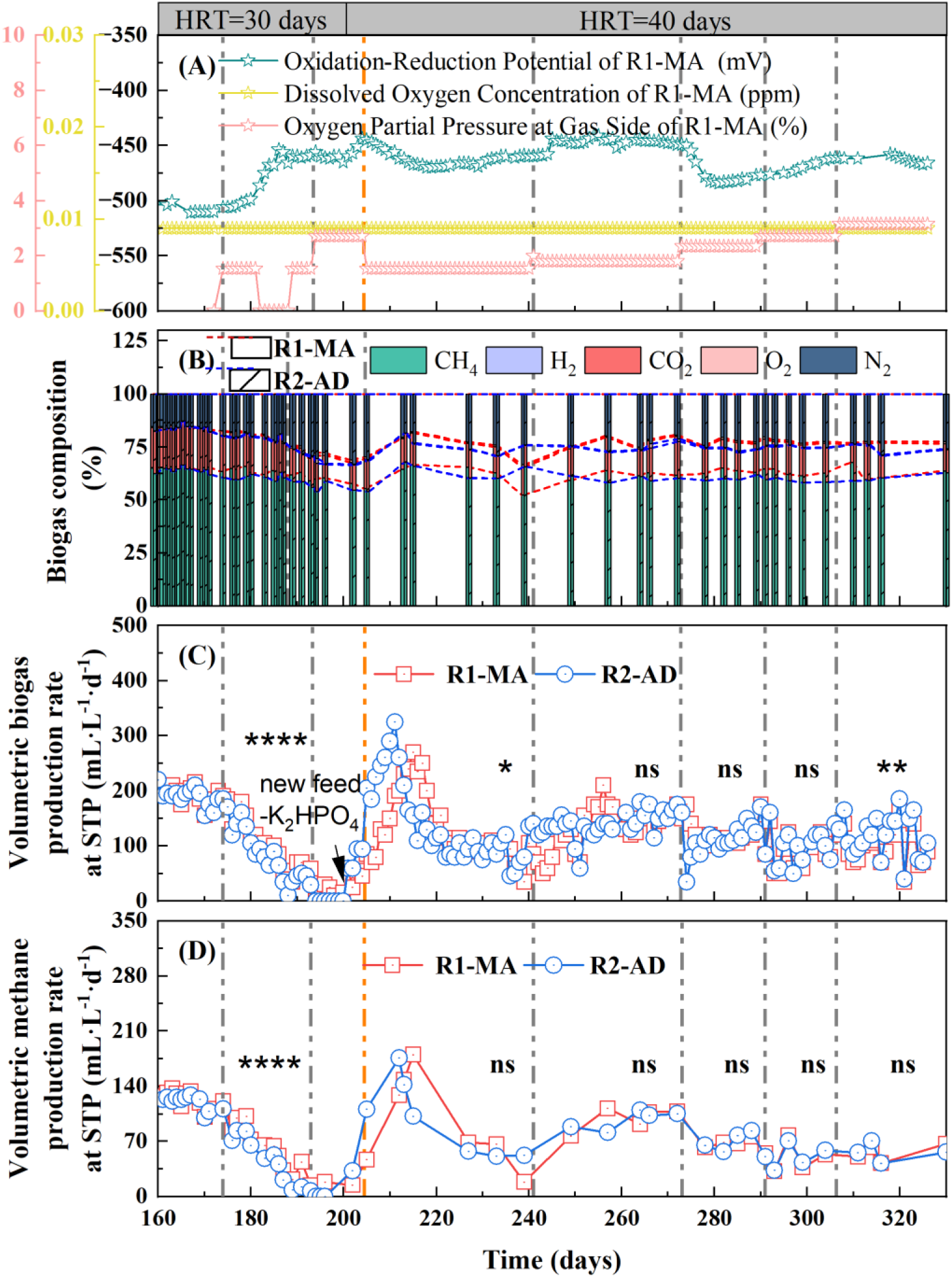
Bioreactor performance: (A) Microaeration intensity given as the oxygen partial pressure on the gas side, oxidation-reduction potential, and dissolved oxygen concentration in bioreactor R1-MA; (B) Biogas composition; (C) Volumetric biogas production rate normalized to 0℃ and 1atm; (D) Volumetric methane production rate. Statistical analysis was conducted using the paired-sample t-tests in Origin. (p-value with asterisks in the figure, ns indicates no significant difference, * indicates p < 0.05, ** indicates p < 0.01, and **** indicates < 0.0001).

The microaeration intensity (*i.e.*, oxygen flux) was changed by adjusting the oxygen partial pressure on the gas side. The correlation coefficient between oxygen flux and oxygen partial pressure was 0.59 based on the experimental design used in the biotic case study (Section 2.2.1). Air at a 1.2 L·h^−1^ flow rate was substituted for the nitrogen flow in the gas flow-through chamber of R1-MA starting on Day 174 to establish an initial oxygen partial pressure of 1.53% with an associated oxygen flux of 0.009 mol·min^−1^ (*i.e.*, 65 mol·L-liquid^− 1^·day^−1^). The oxygen partial pressure was subsequently increased to 1.8%, 1.98%, 2.35%, 2.7%, and 3.15% (**Fig. 5A**), resulting in corresponding oxygen flux values of 76, 84, 99, 115, and 134 mol·L-liquid^−1^·day^−1^, respectively.

The observed inverse correlation between ORP values and oxygen partial pressure highlights the flexibility and precision of controlling microaeration dosing intensities using the membrane bioreactor (**Fig. 5A**). When oxygen partial pressure increases, ORP values rise but with a lag phase of 2 – 6 days (*e.g.*, two days after Day 174, six days after Day 194, and four days after Day 244 in **Fig. 5A**). Although a temporary drop in ORP occurred on Day 274, it continued to increase as the microbiome adjusted to the new microaeration intensity. Conversely, when oxygen partial pressure decreases, ORP tends to increase, suggesting reduced oxidative conditions in response to lower oxygen levels in the system. After Day 214, both oxygen partial pressure and ORP stabilized, with ORP fluctuating slightly but staying relatively constant at a higher level (*i.e.*, – 466 ± 3 mV between Days 214 and 238) than initially recorded (*i.e.*, – 510 ± 2 mV between Days 167 and 174). This similar stabilization pattern also occurred between days 245 and 273 (*i.e.*, – 446 ± 2 mV).

The extended equilibration phase in reaching a stable ORP observed here was most likely caused by the delayed adaptation of the microbiome to the change in oxygen dosing intensity. Also, the ORP did not respond proportionately to the change in oxygen flux. Previous researchers advocate using ORP-controlled oxygen dosing systems for microaeration studies [11, 30]; however, this approach is subject to these complex microbiome responses, leading to irregular and imprecise dosing regimes and poor process control. Microbiomes are unique and dynamic, which suggests the ORP set-points will not always produce the same outcome between bioprocessing applications. These limitations have led to conflicting results between microaeration studies [17]. The membrane bioreactor system presented here provided consistent oxygen gas fluxes independent of biology and the ORP value. Despite this, the biological response to oxygen gas flux was highly varied, suggesting robust bioprocess control in microaeration applications involving open-culture microbiomes may be unobtainable.

#### 3.2.2 Bioreactor performance with microaeration

Between days 175 and 192, microaeration appeared to enhance methane production in R1-MA, even though methane production in both bioreactors declined and eventually ceased by Day 194. Compared to R2-AD, R1-MA maintained 1) significantly higher methane content in the biogas (p < 0.01, Day 175 – 192 in **Fig. 5B**), 2) a higher biogas production rate (p < 0.0001, Day 175 – 192 in **Fig. 5C**), and consequently, 3) a higher methane production rate (p < 0.0001, Day 175 – 192 in **Fig. 5D**). The reduced methane production and carboxylic acid accumulation observed in both bioreactors likely resulted from one or more of the following factors: 1) organic overloading, 2) a short HRT, or 3) inhibition caused by high K_2_HPO_4_ concentrations in the synthetic feed mixture. Notably, when the oxygen partial pressure was increased from 1.53% (O_2_ flux = 9 mmol·min^−1^) to 2.7% (O_2_ flux = 15.9 mmol·min^−1^) on Day 194, biogas and methane production in R1-MA showed a slight recovery after two days, whereas R2-AD remained inactive, producing no methane between days 194 and 200 (**Fig. 5C, 5D**).

Bioprocess instability was characterized by an accumulation of carboxylic acids and high pH (over 8) in both bioreactors (**Fig. 6A, 6B**). Three interventions were implemented to recover bioreactor performance. First, on Day 198, the HRT was increased from 30 to 40 days with OLR decreasing from 1.3 g COD·L^−1^·day^−1^ to 1 g COD·L^−1^·day^−1^, which was followed by a partial reduction in the carboxylic acid concentration (**Fig. 6A**). Due to the elevated pH, the total ammonium concentration was measured to investigate its potential impact. However, the ammonium concentrations in bioreactors R1-MA and R2-AD were 565.5 mg·L⁻¹ and 612.75 mg·L⁻¹ on Day 195, respectively, which remained far lower than the inhibition concentration of 1.7 g·L⁻¹ to methanogenesis [31]. The corresponding free ammonia concentrations, 29.2 and 31.6 mg NH_3–N_·L^−1^ (R1-MA and R2-AD, respectively) were also below the inhibitory concentration of 150 mg NH_3–N_·L^−1^ [24]. Subsequently, K₂HPO₄ was removed from the feed mixture on Day 200 to lower the pH and avoid possible potassium cation inhibition [24, 32], leading to improved methane production in both bioreactors (**Fig. 5D**). Although pH initially decreased from 8.0 on Day 200 to 7.8 on Day 207, it reverted to approximately 8.0 shortly thereafter, suggesting that the elevated pH was not attributable to the K₂HPO₄ buffer in the feed. Lastly, to address the possibility of excessive microaeration intensity, the oxygen partial pressure was reduced to 1.53% (*i.e.*, O_2_ flux = 9 mmol·min^−1^) on Day 205.

**Fig. 6.**
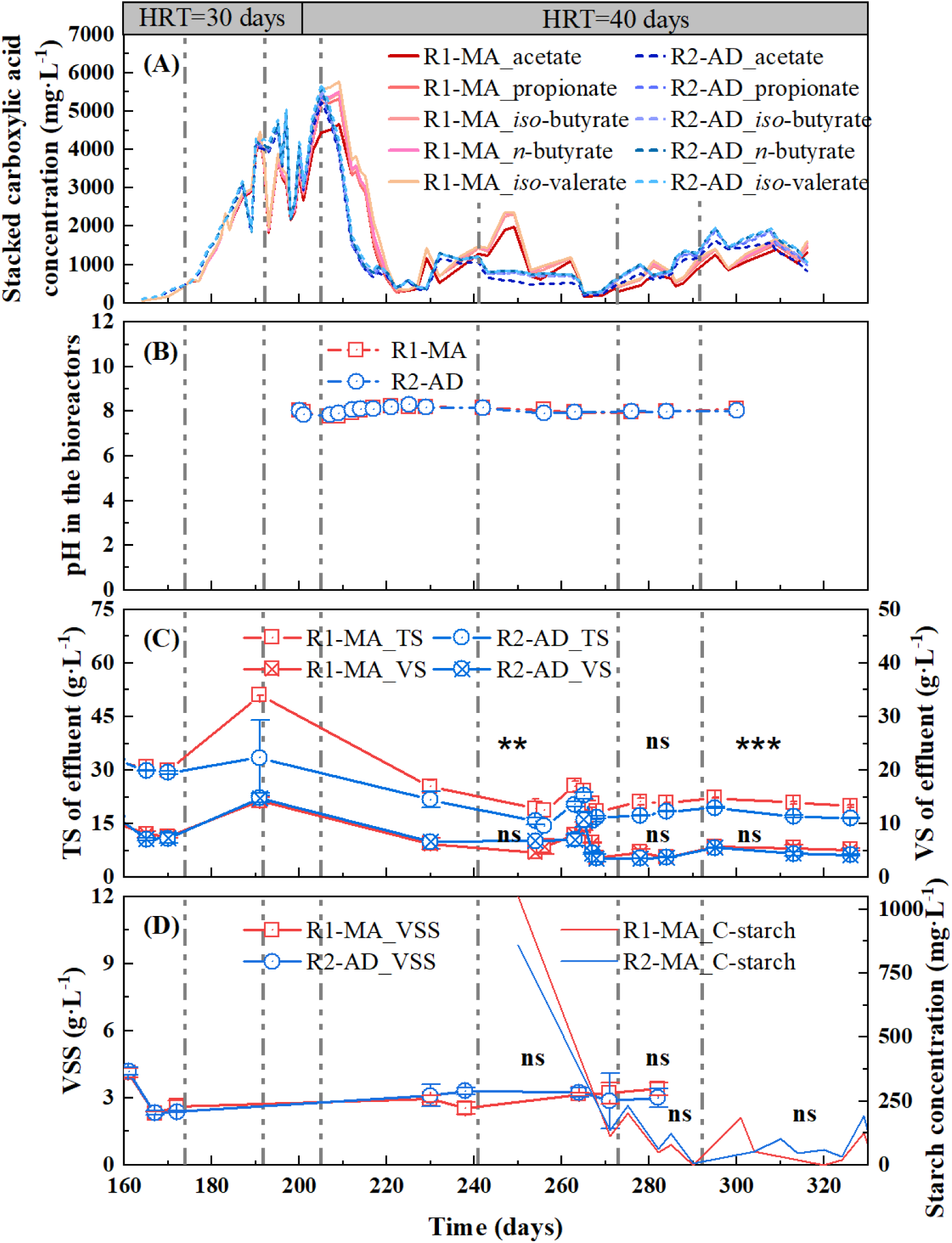
Bioreactor performance characterized by effluent analysis: (A) Carboxylic acid concentration; (B) pH; (C) Total solids (TS) and volatile solids (VS); (D) Volatile suspended solids (VSS) and starch concentrations of the mixed bioreactor broth. Statistical analysis was conducted using the pair-sample t-tests in Origin. (p-value with asterisks in the figure, ns indicate no significant difference, ** indicates p < 0.01, *** indicates p < 0.001).

Following the mentioned interventions, both bioreactors reached a pseudo-steady state between days 222 and 236 with stable biogas and methane production (**Fig. 5C, 5D**), carboxylic acid concentrations lower than 1000 mg·L^−1^ (**Fig. 6A**), and stable pH (**Fig. 6B**). During this period, there was no significant difference in methane production between bioreactors R1-MA and R2-AD (p = 0.08 in **Fig. 5D**). Afterward, we increased the oxygen partial pressure incrementally to 1.8% (*i.e.*, O_2_ flux = 10.6 mmol·min^−1^), 1.98% (*i.e.*, O_2_ flux = 11.7 mmol·min^−1^), 2.35% (*i.e.*, O_2_ flux = 13.8 mmol·min^−1^), 2.7% (*i.e.*, O_2_ flux = 15.9 mmol·min^−1^), and 3.15% (*i.e.*, O_2_ flux = 18.6 mmol·min^−1^) (**Fig. 5A**). Within the range of oxygen microaeration intensity we provided, we did not observe significant improvements in methane production. Although we noticed higher solids concentration in M1-R1 than R2-AD for some time, volatile solids, volatile suspended solids, and starch concentrations were not significantly different between bioreactors (p-value > 0.05) (**Fig. 6C, 6D**). In summary, the microaeration intensities supplied in this study did not show clear enhancement for solids hydrolysis and AD performance, contrary to previous microaeration studies, where this benefit was reported [33–35].

## 4. Concluding remarks

This study successfully designed, characterized, and tested a novel flat-sheet membrane bioreactor for precise trace gas dosing in an experimental-scale bioprocessing application. The key findings of the study include:

(1) Stable mass transfer performance: the integration of mechanical stirring promoted uniform gas distribution, achieving consistent K_L_a values for each membrane. The non-porous silicone membrane exhibited superior long-term stability and resistance to wetting compared to porous membranes, making it ideal for extended operations.
(2) Flexible operational control: adjusting operating parameters such as oxygen partial pressure, gas-side back pressure, membrane type, and total gas flow rate offers flexibility in bioreactor operation, enabling tailored trace gas supply based on specific gas flux requirements.
(3) Consistent gas dosing during long-term bioprocess operation: the biological microaeration case study with starch-containing wastewater validated the system’s effectiveness in supplying consistent gas dosing at trace levels for extended operating periods. Future perspectives: while this study demonstrates the technical viability of the membrane bioreactor system, further work could be performed to 1) optimize bioreactor design for larger-scale experiments, 2) address membrane wetting in porous membrane configurations, 3) modify bioreactor design for high-pressure applications, and 4) explore its application with different gases, substrates, and microbial hosts. Advancing this technology could enable more precise control of gas-dependent bioprocesses and broaden its use in experimental and industrial testing settings. We expect further research and development to result in greater technical performance and higher fidelity.

## Declarations

### Declaration of Generative AI and AI-assisted technologies in the writing process

During the preparation of this work, the first author used ChatGPT to refine the language to achieve a more academic tone. After using this tool, the authors reviewed and edited the content as needed and took full responsibility for the content of the publication.

### Author contributions

Mei Zhou: Conceptualization, Investigation, Visualization, Writing - Original Draft. Ibrahim Bello: Investigation, Visualization, Writing - Original Draft of CFD. Jose Antonio Magdalena: Conceptualization, Investigation. Joseph G. Usack: Supervision, Conceptualization, Investigation, Writing - Review & Editing, Funding acquisition.

### Funding and acknowledgments

This work was supported, in part, by grants from 1) the German-Israeli Foundation for Scientific Research and Development (GIF) (grant number I-1547-500.15/2021), 2) the University of Georgia’s Institute of Integrative Precision Agriculture (IIPA) Seed Grant program, and 3) the University of Georgia’s Agricultural Experiment Station. The authors would like to acknowledge Dr. Gerben Stouten for supporting data verification.

