## Supplementary Information for "Design and validation of a new flat-sheet membrane bioreactor system for bioprocessing research requiring very low gas fluxes"

**[*]Corresponding author:**

### Supplementary Information

#### SI l: Computational Fluid Dynamic (CFD) Simulation

The CFD simulation results are presented as a 2D cross-section of the bioreactor indicating the fluid velocity (m·s$^{-1}$) using a color contour plot for three mixing speeds (*i.e.*, 250 rpm, 350 rpm, 450 rpm) (**Fig. S1**). Fluid velocity was reasonably uniform and symmetric throughout the bioreactor volume for each simulated mixing speed, with higher fluid velocities occurring near the stir bar and the bioreactor body wall and lower fluid velocities occurring in the central fluid column. The CFD simulation indicated turbulent mixing conditions throughout the bioreactor volume with no significant dead zones. Moreover, because the fluid packets entrain the dissolved oxygen molecules, we can assume dissolved oxygen dynamics mimic the fluid velocity behavior within the bioreactor under abiotic conditions. These results also suggest that the ORP and DO sensor readings represent the bulk concentration and that the mechanical stirring system near the membrane surface is sufficient to prevent dissolved oxygen stratification.

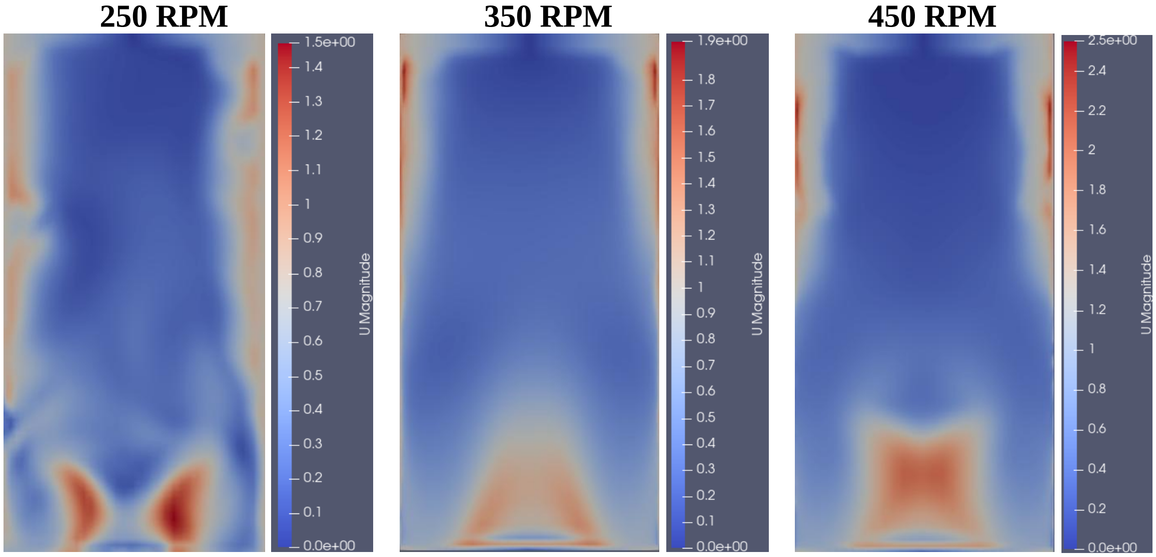


**Fig. S1** Computational fluid dynamic simulation of the bioreactor as a 2D cross-section showing the fluid velocity (m·s$^{-1}$) at three mixing intensities: 250 rpm, 350 rpm, and 450 rpm. Blue coloring indicates a lower fluid velocity, while red coloring indicates a higher fluid velocity.

In OpenFOAM, the following equations were applied in the CFD analysis based on the volume of fluid (VoF) method.

$$\frac{\partial\alpha}{\partial t} + V.\nabla\alpha + \nabla.(C_\alpha|V|\alpha(1 - \alpha)\frac{\nabla\alpha}{|\nabla\alpha|}) = 0 \quad [1]$$

$$\rho = \rho_1\alpha + \rho_2(1 - \alpha) \quad [2]$$

$$V = V_1\alpha + V_2(1 - \alpha) \quad [3]$$

$$\mu = \mu_1\alpha + \mu_2(1 - \alpha) \quad [4]$$

*where:*

$\alpha$ = the volume fraction

$C_\alpha$ = the artificial compression coefficient

$t$ = the time, s

$\rho$ = density of the mixture, kg·m$^{-3}$

$\rho_1$ = density of sludge, kg·m$^{-3}$

$\rho_2$ = density of the void (air), kg·m$^{-3}$

$V$ = velocity of the mixture, m·s$^{-1}$

$V_1$ = velocity of sludge, m·s$^{-1}$

$V_2$ = velocity of air, m·s$^{-1}$

$\nabla$ = the spatial gradient operator, m$^{-1}$

$\mu$ = dynamic viscosity of the mixture, Pa·s

$\mu_1$ = dynamic viscosity of sludge, Pa·s

$\mu_2$ = dynamic viscosity of air, Pa·s

The continuity and Navier-Stokes equations were used to describe the physics of fluid flow:

**Conservation of Mass**

$$\frac{\partial\rho}{\partial t} + \nabla.(\rho V) = 0 \quad [5]$$

The above equation simplifies to:

$$\frac{\partial\rho}{\partial t} + \frac{\partial\rho u}{\partial x} + \frac{\partial\rho v}{\partial y} + \frac{\partial\rho w}{\partial z} = 0 \quad [6]$$

**Conservation of Linear Momentum**

$$\frac{\rho\partial V}{\partial t} + \rho V.\nabla(V) = \nabla.[-pI + K] + \rho g + f \quad [7]$$

$$f = F_s + F_B \quad [8]$$

*where*:

$P$ = the pressure, Pa

$I$ = the identity matrix

$K$ = the viscous stress tensor, Pa

$G$ = the gravitational acceleration vector, m·s$^{-2}$

$\mu_T$ = turbulent viscosity, Pa·s

$\tau_{xz}$ = the shear stress component, Pa

$V_x$, $V_y$, $V_z$ = the velocity components in x/y/z, m·s$^{-1}$

$\partial/\partial x$, $\partial/\partial y$, $\partial/\partial z$ = the derivatives in x/y/z directions, m$^{-1}$

$f$ = the total force per unit volume, N·m$^{-3}$

$F_s$ = the surface forces, N·m$^{-3}$

$F_B$ = the body forces, N·m$^{-3}$

Expressing [7] in cartesian coordinates, we have:

***x - component***

$$\frac{\rho \partial V_x}{\partial t} + \frac{\rho V_x \partial V_x}{\partial x} + \frac{\rho V_y \partial V_x}{\partial y} + \frac{\rho V_z \partial V_x}{\partial z} = -\frac{\partial p}{\partial x} + (\mu + \mu_T)\left(\frac{\tau_{xy} + \tau_{xz}}{\mu}\right) + f_x \quad [9]$$

***y - component***

$$\frac{\rho \partial V_y}{\partial t} + \frac{\rho V_x \partial V_y}{\partial x} + \frac{\rho V_y \partial V_y}{\partial y} + \frac{\rho V_z \partial V_y}{\partial z} = -\frac{\partial p}{\partial y} + (\mu + \mu_T)\left(\frac{\tau_{xy} + \tau_{yz}}{\mu}\right) + f_y \quad [10]$$

***z - component***

$$\frac{\rho \partial V_z}{\partial t} + \frac{\rho V_x \partial V_z}{\partial x} + \frac{\rho V_y \partial V_z}{\partial y} + \frac{\rho V_z \partial V_z}{\partial z} = -\frac{\partial p}{\partial z} + (\mu + \mu_T)\left(\frac{\tau_{xz} + \tau_{yz}}{\mu}\right) + \rho g + f_z \qquad [11]$$

#### SI II: Startup Period of the Biotic Microaeration Study

The inoculum used to inoculate the membrane bioreactors for the biotic microaeration study was collected from an active 5.5 L laboratory anaerobic digester, producing 558 mL $CH_4 \cdot L^{-1} \cdot day^{-1}$ at a pseudo-steady state. The substrate mixture fed to the membrane bioreactor was composed of (per liter): 13.64 g potato starch, 13.64 g sodium acetate, 4.09 mL propionic acid, 3.41 g casein, 0.1 g yeast extract, 5 g sodium sulfate, 10 g di-potassium hydrogen phosphate, 30 mL mineral stock, and 30 mL trace element solution. The mineral stock contains the following (in g/L): $NaCl_2$, 40; $NH_4Cl$, 50; KCl, 5; $KH_2PO_4$, 6; $MgCl_2$, 7; $CaCl_2$, 2. The trace element solution consists of (in g/L): $C_6H_9NO_6$, 2; $MnSO_4$,1; $FeSO_4$, 0.8; $CoCl_2$, 0.2; $ZnSO_4$, 0.2; $CuCl_2$, 0.02; $NiCl_2$, 0.02; $Na_2MoO_4 * 2H_2O$, 0.02; $Na_2SeO_4$, 0.02; $Na_2WO_4$, 0.02. During the experimental phase, the organic loading rate (OLR) changed from 2, 1.3, and 1 g $COD \cdot L^{-1} \cdot day^{-1}$, corresponding to the HRT of 20, 30, and 40 days, respectively, to stabilize the bioprocess.

During the startup period, several operational parameters were optimized to enable the bioreactors to reach pseudo-steady state operation. Here, we increased the hydraulic retention time (HRT) from 20 to 30 days to increase biomass retention. We modified the synthetic feed mixture by removing sodium sulfate ($Na_2SO_4$) on Day 110, which increased the biogas production rates (**Fig. S2A**), methane content in the biogas **(Fig. S2B)**, and methane production rates (**Fig. S2C)**. Biogas performance reached a pseudo-steady state during Day 134 – 154 (**Fig. S2**). A carboxylic acid accumulation issue was also resolved after removing $Na_2SO_4$ (**Fig. S3A**). To eliminate the slight difference in the microbial community and solids concentration in both bioreactors, we mixed the broth in the bioreactors on Day 155 and reinoculated them. The ORP probe was recalibrated on the same day (**Fig. S2D)**. After mixing the bioreactors, there was still a slight difference in biogas production due to a leak in bioreactor R2-AD (Day 156 – 159 in **Fig. S2A**), which was fixed on Day 159. Chemical oxygen demand (COD, **Fig. S3B**), total solids and volatile solids (TS and VS, **Fig. S3C**), and volatile suspended solids (VSS, **Fig. S3D**) behaved equally between bioreactors ($p < 0.05$).

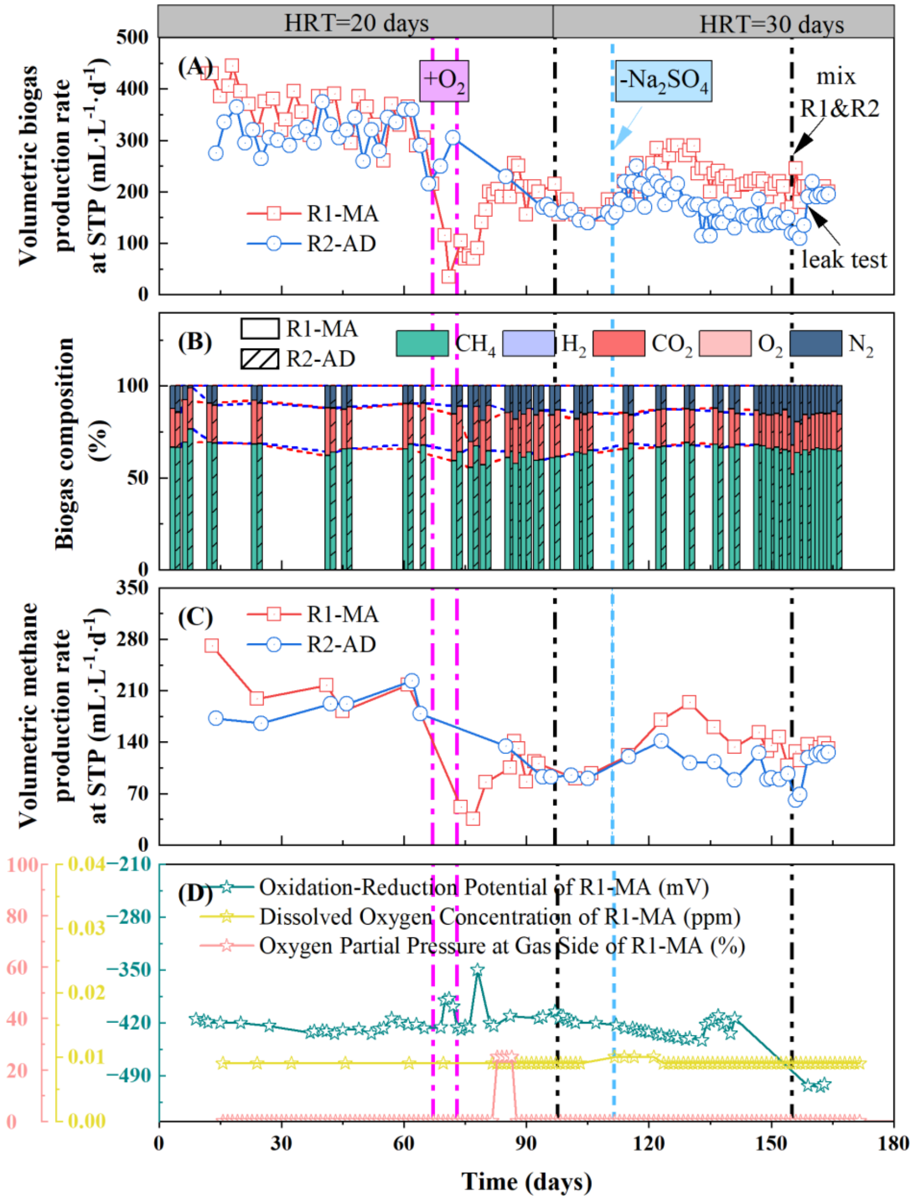


**Fig. S2** Startup phase before Day 155: (A) Volumetric biogas production rate normalized to 0℃ and 1 atm conditions; (B) Biogas composition; (C) Volumetric methane production rate normalized to 0℃ and 1atm conditions; (D) Microaeration intensity as the oxygen partial pressure in the gas side, oxidation-reduction potential, and dissolved oxygen concentration in bioreactor R1-MA.

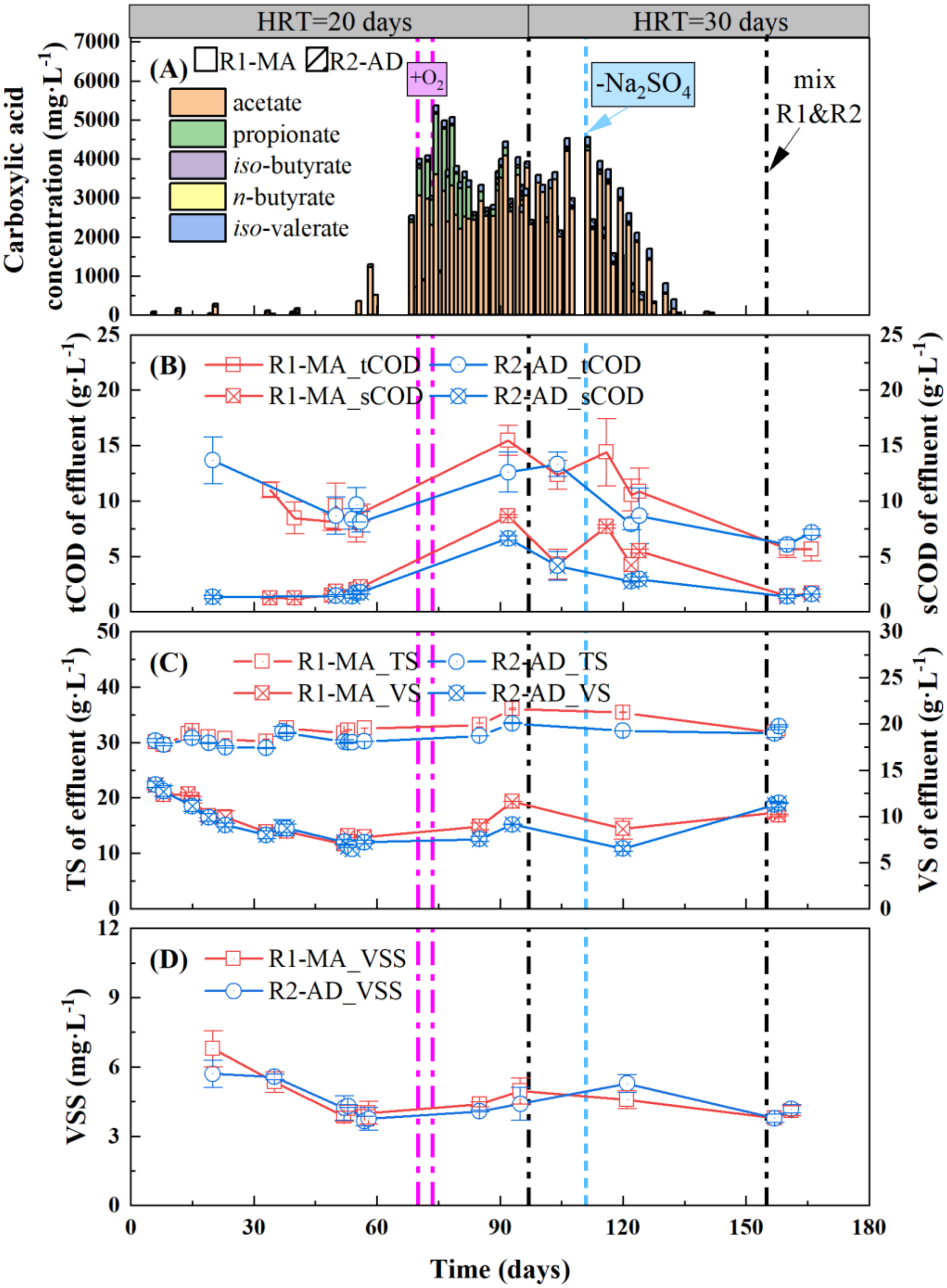


**Fig. S3** Bioreactor performance characterized by effluent analysis during the startup phase before Day 155: (A) Carboxylic acid concentration; (B) Chemical oxygen demand (COD); (C) Total solids (TS) and volatile solids (VS); (D) Volatile suspended solids (VSS) of mixed bioreactor broth.
